# Genomic instability and biofilm determinants in *Streptococcus mutans*: insights from a sequence-defined arrayed transposon library

**DOI:** 10.64898/2026.03.25.714184

**Authors:** Ana Karen Solano Morales, Emanuel Cazano, Cara Pirani, Graysen Jones, Andrew Goode, Alejandro R. Walker, Anthony Sperduto, Bikash Dwivedi, Pallavi Bantha, Stella D. Peter, Lisa K. McLellan, Mohamed A. Alam, Robert C. Shields

## Abstract

*Streptococcus mutans* is a primary architect of dental caries, utilizing complex genetic networks to build resilient, acid-producing biofilms. While pooled screens (Tn-seq) have identified important fitness factors, they often fail to capture extracellular or moderate-effect determinants due to community-level masking. Therefore, to study biofilm phenotypes, we constructed a comprehensive arrayed library of 9,216 mutants and used Cartesian Pooling-Coordinate Sequencing (CP-CSeq) to establish a sequence-defined resource covering 51% of non-essential genes. By screening the entire collection in isolation, we identified several novel biofilm determinants, including the putative metal transporter SMU_635 and the glycosylation-associated protein SMU_2160. However, systematic whole-genome sequencing (WGS) of our hits revealed an interesting level of genomic instability: 25% of biofilm-defective mutants had undergone spontaneous recombination at the *gtfBC* locus, while 7% had lost the Tn*Smu1* element, an excision rate 1,000-fold higher than previously reported. While targeted mutagenesis confirmed that Tn*Smu1* loss does not impact biofilm integrity, the *gtfBC* deletions directly accounted for the most severe phenotypes, highlighting a systemic risk of misattributing gene functions to primary transposon insertions. Our findings provide a powerful new genetic resource for the *S. mutans* community while establishing a critical new standard: an arrayed library is only as defined as its underlying genome, making systematic genomic verification an essential prerequisite for accurate functional genomics.

**Importance:** *Streptococcus mutans* is a major human pathogen responsible for dental caries, a global public health challenge driven in part by the organism’s ability to form resilient, acidogenic biofilms. While traditional pooled genetic screens have identified many fitness factors, they often fail to capture extracellular or moderate-effect determinants because neighboring healthy bacteria can mask these defects. This work provides the scientific community with a sequence-defined arrayed mutant library, an essential resource for dissecting the individual contributions of genes to biofilm integrity in isolation. Beyond identifying well-known machinery, this study uncovers novel determinants, including the putative metal transporter SMU_635 and the putative glycosylation-associated protein SMU_2160. Crucially, the discovery of pervasive genomic instability within the library, specifically at the *gtfBC* and Tn*Smu1* loci, reveals a systemic risk in functional genomics: the potential to misattribute phenotypes to primary mutations when the underlying background has undergone large-scale rearrangements. By establishing systematic whole-genome verification as a necessary standard, this research ensures that the identification of future therapeutic targets is built upon a verified genetic foundation.

## Introduction

*Streptococcus mutans* is recognized as the primary etiological agent of dental caries due to its exceptional capacity to both survive in acidic environments and form robust, acid-producing biofilms (1). Biofilm development begins with bacterial adhesion to the salivary pellicle, mediated by both sucrose-dependent and sucrose-independent pathways. The sucrose-dependent mechanism relies heavily on glucosyltransferases (Gtfs B, C, and D), which synthesize extracellular glucans that are critical for cell adhesion and the structural integrity of the extracellular matrix (2). Further strengthening of the biofilm is provided by glucan-binding proteins (Gbps), which promote bacterial aggregation and biofilm cohesiveness (3, 4). Sucrose-independent adhesion is mediated by high-affinity surface proteins, such as Antigen I/II (P1, SpaP), which facilitate colonization and coaggregation with other oral bacteria (5–7). These complex genetic mechanisms enable *S. mutans* to establish a resilient and multispecies biofilm characteristic of the carious lesion. Despite significant knowledge of the genetic determinants of *S. mutans* biofilm development, because this is a crucial step in dental caries development, it remains essential to systematically identify all genetic factors required for robust biofilm formation. Additionally, due to the extensive characterization of *S. mutans* biofilm formation, this process serves as an ideal benchmark for validating new functional genomics platforms, as a robust system should reliably identify well-established determinants as a prerequisite for the discovery of novel ones.

Unbiased functional genomic approaches are indispensable for dissecting the complex genetic networks governing *S. mutans* physiology. While transposon insertion sequencing (TIS), including methods like Tn-seq, has become a standard tool for assessing genome-wide fitness contributions and identifying essential genes (8), these pooled approaches have a critical limitation. The reliance of TIS on pooled, competitive growth, presents a significant barrier to studying community-dependent traits. In a mixed population, defects in extracellular traits, such as biofilm matrix production, can be masked by neighboring cells expressing wild-type levels of these critical biofilm factors. Arrayed mutant libraries resolve this by facilitating direct genotype-to-phenotype mapping through systematic, high-throughput monoculture screening, a strategy that has revolutionized functional genomics in model organisms like *E. coli* (9). Leveraging recent advances in sequence-based deconvolution (10–13), which allow for the creation of arrayed libraries from pooled TIS inputs, we aim to generate an arrayed transposon mutant library for *S. mutans* using an existing pooled Tn-seq library (8). This approach allows for the high-throughput identification of biofilm-defective mutants, offering new insights into the genetics of tooth decay that pooled methods may overlook.

That being said, a significant, yet often overlooked, challenge in high-throughput functional genomics is the inherent genomic stability of the target organism. In *S. mutans*, repetitive sequences and mobile genetic elements (MGEs) may represent potential hotspots for large-scale rearrangements that can occur during the multi-step process of library construction, such as transformation, selection on agar, and long-term storage. Because these events can occur at frequencies orders of magnitude higher than typical point mutations, they pose a risk to data integrity. If these events go undetected, they can lead to misattribution of phenotypes to specific transposon insertions, thereby compromising the reproducibility of the screen and inflating the false-discovery rate. Consequently, the genomic identity of an arrayed library is only as reliable as the stability of its underlying background genome. In this study, we demonstrate that incorporating systematic whole-genome verification is not merely a validation step, but a critical requirement for differentiating true gene-phenotype links from high-frequency genomic artifacts.

In this study, we apply a deconvolution strategy, based on Cartesian Pooling-Coordinate Sequencing (CP-CSeq) (11), to create a sequence-defined *S. mutans* arrayed transposon mutant library. We systematically characterized this resource to identify genetic determinants of sucrose-dependent biofilm formation. Crucially, we combined high-throughput screening with systematic whole-genome sequencing and PCR-based verification to assess the genomic integrity of the library. This approach allowed us to quantify the frequency of high-frequency recombination at the *gtfBC* locus and the spontaneous excision of the integrative and conjugative element (ICE) Tn*Smu1*. Through this dual focus, our study provides the *S. mutans* community with a sequence-defined, arrayed genetic resource while establishing a critical methodological framework for differentiating true gene-phenotype links from high-frequency genomic artifacts. Our results provide new insights into the requirements for biofilm formation and highlight the essential role of genomic verification in streptococcal functional genomics.

## Results

### Arrayed mutant library construction and deconvolution

To establish a resource for functional genomics, 9,216 single *S. mutans* transposon mutants were isolated from a previously generated library (8) and processed via several steps (Figure 1). First, mutants were arrayed into 96-well plates containing BHI-spectinomycin media, with each plate assigned a unique identifier (e.g., Tn-SmUA159-P1). The entire arrayed library was stored in triplicate and stored at –80 °C, providing a resource for high-throughput phenotypic screening. Next, the genomic location of the Tn insertion for each mutant was determined using combinatorial pooling followed by CP-CSeq deconvolution (11). This approach greatly reduces the complexity and cost of sequencing. In brief, mutants were pooled from the arrayed plates to create 40 samples, rather than 9,216 individual samples, ensuring each insertion was represented in four distinct groups for accurate deconvolution. The pooling scheme included: X-pools, where all mutants in the same row were combined; Y-pools, where all mutants in the same column were combined; and Z-pools, where mutants from each 96-well plate were combined into a single well of a 96-well plate and then row and column pools were generated from this master plate. These pools allow the unique identification of each mutant based on its combinatorial signature across the four X, Y, and Z (row and column) pools (Figure S1).

**Figure 1.**
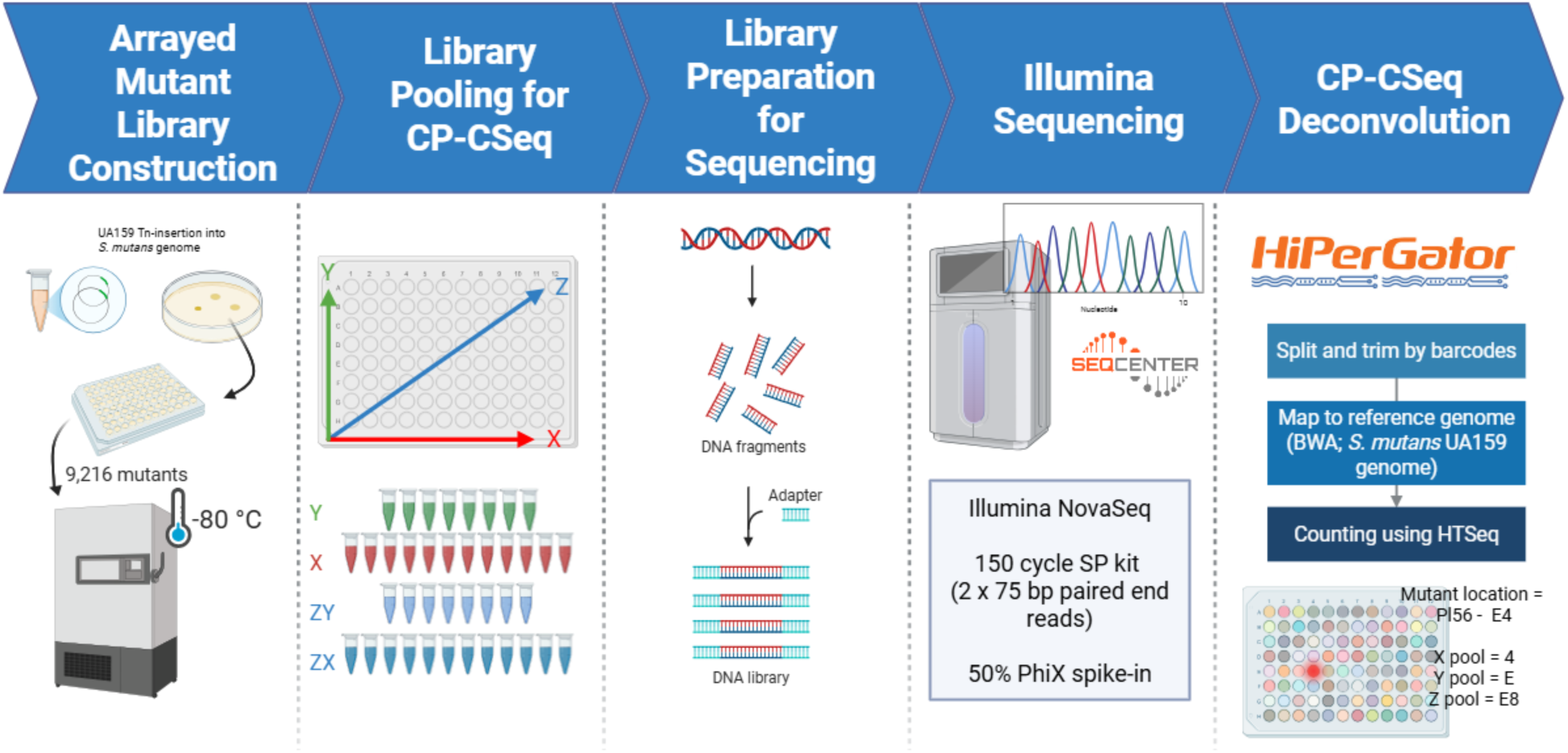
Overview of the arrayed mutant library construction, sequencing, and deconvolution workflow. The systematic generation of the sequence-defined *S. mutans* arrayed library was achieved through a multi-step pipeline: (1) Single transposon mutants were isolated from a pooled *S. mutans* UA159 library and inoculated into 96-well plates to create a physical resource for high-throughput screening, (2) to facilitate cost-effective sequencing, mutants were combined into 40 distinct combinatorial pools (X, Y, and Z axes) such that each individual mutant’s genomic coordinates could be identified by its unique presence across four specific pools, (3) genomic DNA was extracted from the pooled samples, followed by enzymatic digestion and adapter ligation to prepare DNA fragments for high-throughput sequencing, (4) the prepared libraries were sequenced using the Illumina NovaSeq platform to capture the flanking regions of each transposon insertion, and (4) raw sequence data were processed using high-performance computing (HiPerGator) to map reads back to the *S. mutans* reference genome and decode the physical plate coordinates for each identified insertion site.

The pooled libraries were sequenced using the Illumina NovaSeq platform. Sequence data processing was performed using the University of Florida HiPerGator high-performance computer cluster. Reads were 1) demultiplexed by barcode and the 16 bp downstream of the Tn insertion site were extracted, 2) aligned to the *S. mutans* UA159 reference genome using BWA, and 3) quantified using HTSeq after being mapped. A total of 2,498,303,498 reads were obtained from the Illumina NovaSeq runs. After trimming, 2,105,987,502 (84.3%) reads were obtained and 1,861,056,846 were mapped to the *S. mutans* UA159 reference genome, yielding an unmapped rate of 4.75%. The average depth was 47,793,946 reads per barcode. While the previously published CP-CSeq pipeline (11) was useful for designing the sequencing strategy for the transposon mutant population, custom computational scripts were developed to deconvolve the pooled sequencing results (see Supplementary Materials). Of the 9,216 mutants 1,400 mutants were successfully deconvoluted by appearing in four combinatorial pools, confirming high-confidence assignment (Figure 2A). We also assigned a total of 431 mutants where reads appeared in two pools at specific coordinates (e.g. X) but there was a much larger read count from one of the pools. If reads mapped to <4 pools or mapped ambiguously to multiple genomic positions, they were discarded. After our analysis, a total of 894 distinct genes were identified as containing at least one transposon insertion. This saturation resulted in coverage of 51% of the total non-essential protein-coding genes in *S. mutans* UA159 (Figure 2B). The transposon insertions were randomly and densely distributed, with an average distance between insertion sites of ∼1,107 bp, confirming a highly saturated library (Figure 2C). Analysis of intragenic insertion sites revealed no significant positional bias, with insertions distributed uniformly across the normalized lengths of the targeted open reading frames (Figure 2D). The entire collection of deconvoluted transposon mutants is shown in Table S1.

**Figure 2.**
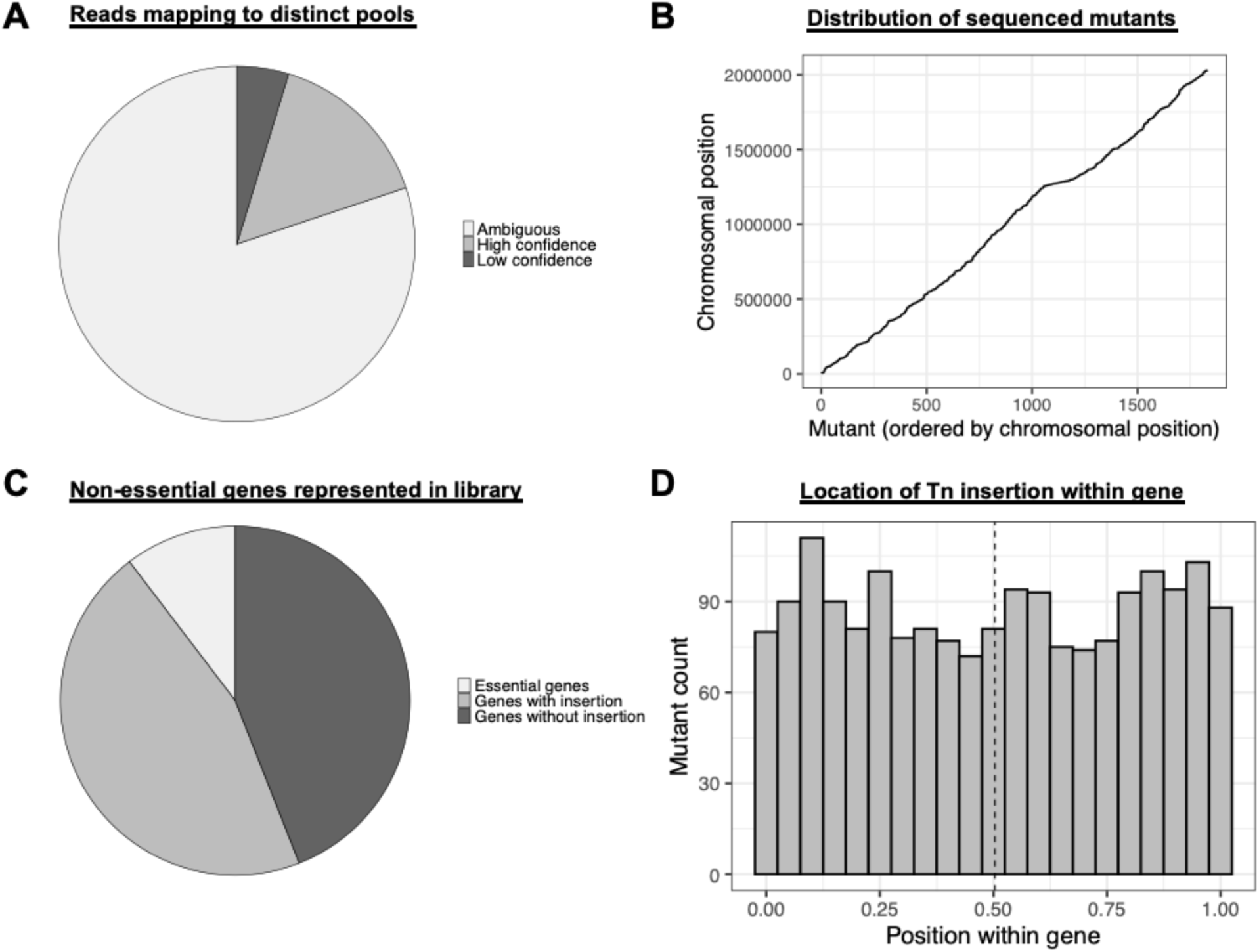
Deconvolution metrics and genomic distribution of the arrayed transposon library. (A) High-confidence assignment was achieved for 1,400 mutants that appeared in four distinct combinatorial pools. Ambiguous reads, including those mapping to fewer than four pools, were discarded to maintain data integrity. (B) Successfully deconvoluted mutants are plotted by their chromosomal position against their order in the library. The linear distribution confirms that transposon insertions are randomly and densely distributed across the *S. mutans* UA159 genome. (C) The library contains insertions in 51% of the non-essential protein-coding genes in the *S. mutans* UA159 reference genome. (D) The histogram displays the location of transposon insertions within the open reading frames (ORFs) of targeted genes. The relatively even distribution across the length of the genes (0.0 to 1.0) indicates a lack of significant insertion bias toward the 5’ or 3’ ends.

To assess the accuracy of CP-CSeq deconvolution, twenty mutants were randomly selected for PCR verification. Primers were designed that would amplify approximately 500 bp up- and downstream of the expected transposon insertion. The expected transposon insertion was confirmed for 16 out of 20 mutants (80%) (Figure S2). These strains produced amplicons consistent with CP-CSeq predicted insertion sites as the resulting PCR amplicon was ∼1,100 bp larger than in the wild-type strain (which is the size of the transposon cassette).

### Demonstrating library utility with a biofilm screen

While CP-CSeq successfully mapped a core set of ∼1,800 mutants with high confidence, the physical array of 9,216 strains provides a much broader landscape for phenotypic discovery. With this in mind, we next evaluated the functional utility of the collection, performing a comprehensive screen of the entire 9,216-mutant library for sucrose-mediated biofilm formation. A key advantage of the arrayed format is the ability to conduct high-throughput phenotypic assays across the entire population, including the ∼80% of mutants that were not deconvoluted. We systematically screened the 9,216 arrayed transposon mutants for defects in sucrose-dependent biofilm formation using a crystal violet assay. To quantify the phenotypic impact of each mutation, we normalized the biofilm biomass from three biological replicates into a biofilm score (Z-score). Across the entire screened collection, the average biofilm score was near zero (−0.008 ± 0.009 S.E.) with a median of 0.06, indicating that the majority of mutants retained a wild-type biofilm phenotype. Using a strict cut-off, a total of 687 mutants were classified as biofilm-defective (biofilm score < −1) and 538 were classified as biofilm-overproducers (biofilm score > +1). The distribution of the biofilm scores across the library is displayed in Figure 3.

**Figure 3.**
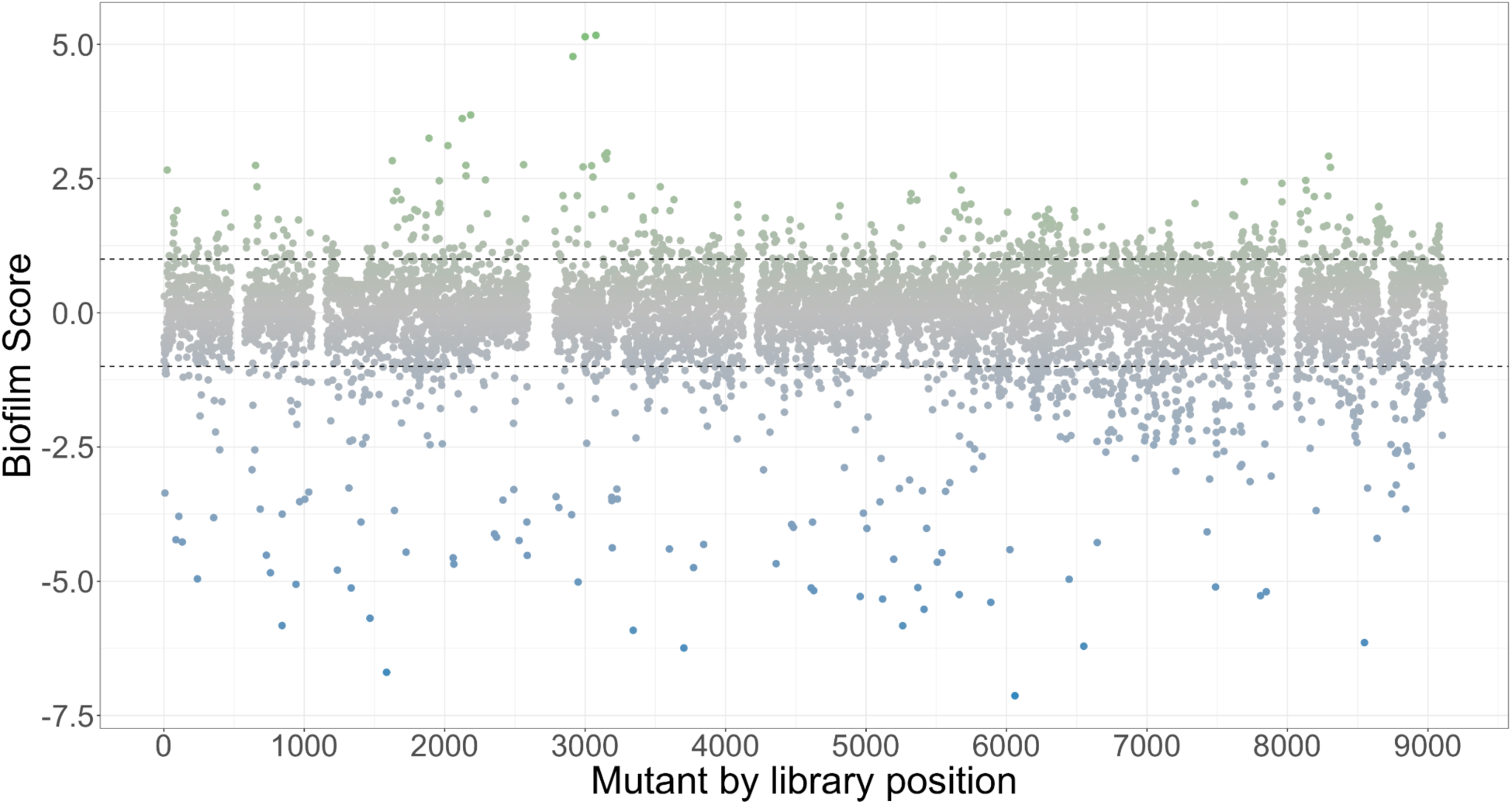
Distribution of biofilm-deficient phenotypes across the arrayed transposon library. The relative biofilm formation of ∼9,216 arrayed mutants was assessed via sucrose-mediated crystal violet assays and expressed as Z-scores. Strains with a biofilm score (Z-score) < −1 were classified as biofilm-defective. A distinct tail of severe outliers (biofilm score < −2.5) identifies the most critical determinants, while most of the library exhibits a wild-type phenotype between a biofilm score of 1 and −1.

To identify the genetic basis for the observed biofilm defects, we selected 84 of the 687 mutants with a biofilm score < −1 for genomic characterization. Rather than selecting only mutants with the most severe phenotypes (e.g., biofilm score < −4), we intentionally sampled strains across the entire defective range. This strategy was designed to increase the probability of identifying novel genes with moderate phenotypes, as well as potential hypomorphic alleles (e.g., 3’ transposon insertions or insertions in promoter regions) that might not produce as strong a phenotype as a full gene knockout. The biofilm scores and genetic backgrounds for these 81 mutants are shown in Figure 4, with comprehensive genomic details, including insertion locations, SNPs, and chromosomal rearrangements, provided in Tables S2 and S4. Analysis of this 81-mutant subset confirmed that, as anticipated, the most severe biofilm defects (scores < −2.5) were dominated by mutations in genes with established, major roles in sucrose-mediated biofilm formation. A total of 59% of mutants in this strong-defect category had insertions affecting *gtfBC* or their regulator, *gcrR*. Notably, *gcrR* mutants had insertions in the upstream intergenic region (between *gcrR* and SMU_1925c), which likely exert polar effects on *gcrR* expression. In contrast, the mutants with moderate biofilm defects (biofilm score between - 1 and −2.5) displayed significantly greater genetic diversity. In this group, only 9% of the strains were *gtfBC* or *gcrR* mutants. The remaining mutants included insertions in numerous genes previously linked to biofilm formation or carbohydrate utilization, such as *scrB*, *manZ*, *mntH*, *lacD2*, *brpA*, *mubM*, *rgpI*, *fruR*, *fruI*, *ciaH*, *pgfM1*, SMU_63c, and *copA*. Beyond the well-characterized determinants of biofilm formation, the screen identified several genes with no previously described role in *S. mutans* sucrose-dependent biofilm development. Among the most significant novel hits was an insertion in an HNH endonuclease signature motif-containing protein (SMU_47, Z-score = −5.83) and a cluster encoding inner membrane and ATP-binding proteins (SMU_429c–SMU_431, Z-score = −4.5). Genes involved in fundamental cellular maintenance and nutrient acquisition were also identified as necessary for optimal biofilm growth, including the glycerol dehydrogenase *gldA* (SMU_495, Z-score = −1.45), the putative dihydrolipoamide acetyltransferase *adhC* (SMU_129, Z-score = −1.94), components of the ABC transporter system responsible for importing the amino acid histidine (*hisJ*, SMU_242c, Z-score = −3.33; *hisM*, SMU_1521, Z-score = −1.2), and the putative dihydroxy-acid dehydratase *ilvD* (SMU_2128, Z-score = −3.73). The identification of these diverse functional categories suggests that sucrose-mediated biofilm formation is integrated into the broader physiological and metabolic state of the cell, extending well beyond the primary extracellular matrix production machinery.

**Figure 4.**
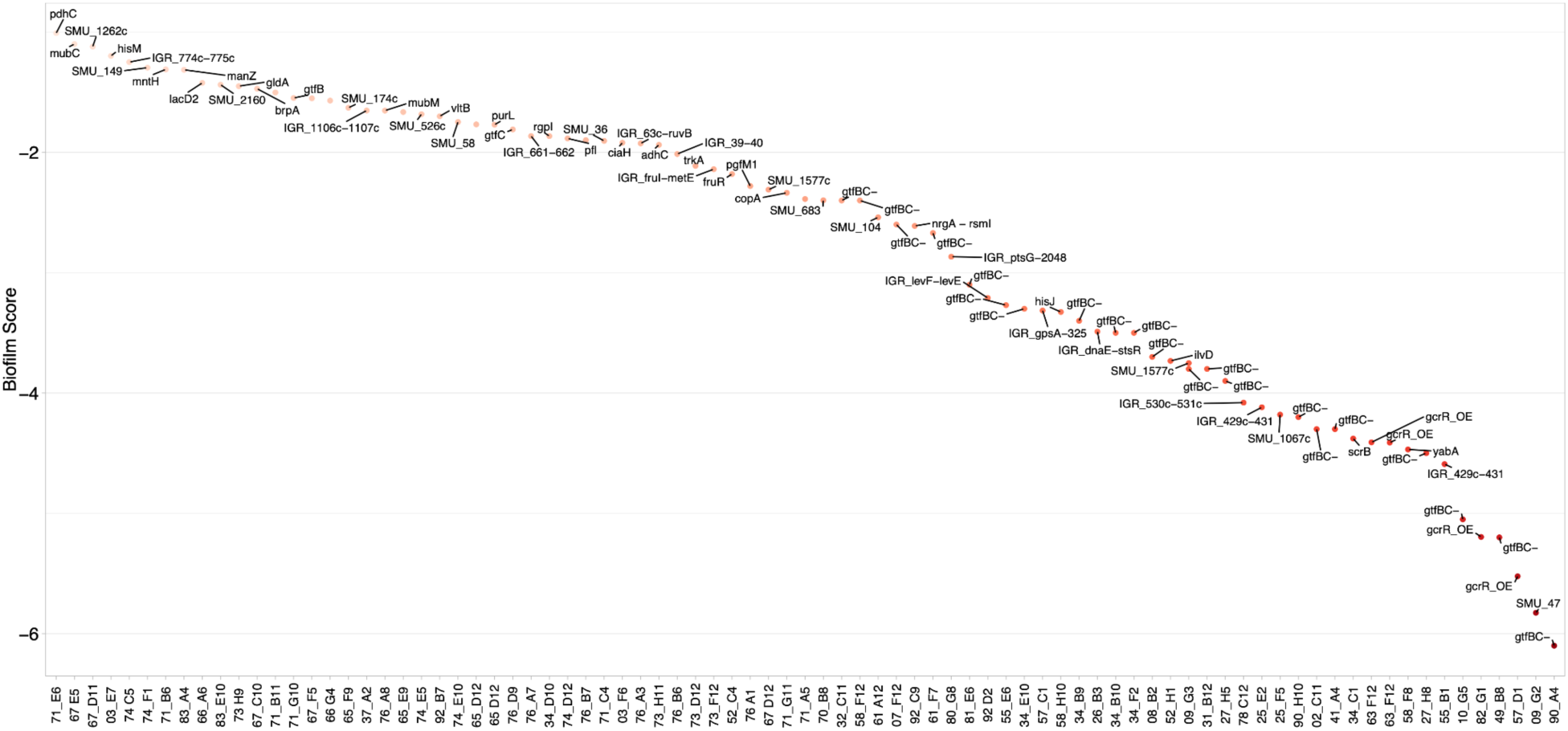
Biofilm phenotypes of whole-genome sequenced transposon mutants. Biofilm formation Z-scores (y-axis) are plotted for each individual mutant (x-axis) selected for comprehensive genomic characterization. Each data point represents a specific mutant whose genomic identity was established via whole-genome sequencing to differentiate between primary transposon-mediated defects and those arising from spontaneous large-scale rearrangements. The distribution illustrates the phenotypic range of the sequenced subset, highlighting that the most severe biofilm-deficient phenotypes predominantly correlate with deletions at the *gtfBC* locus. Each of these mutants is described in more detail in Table S2.

### Genome sequencing identifies areas of gene loss in transposon mutants

To ensure the most accurate interpretation of the data, we genome sequenced a total of 84 biofilm-defective mutant strains. As bacterial whole-genome sequencing costs have fallen it has become more attainable to sequence larger numbers of mutant strains. In the context of transposon mutant screens, genome sequencing helps with identifying factors that are causing mutant phenotypes that are not related to insertion within a gene. These could include genome rearrangements, off-target mutations, and polar effects that disrupt downstream genes. To our knowledge, a systematic verification of transposon mutant backgrounds has not been conducted for this type of screen before, and yet, has clear utility for increasing the reproducibility and robustness of transposon mutant screens.

Initial genome sequencing of a limited number of mutants revealed spontaneous recombination events within the highly homologous *gtfBC* region. These two genes share significant similarity (88.6% identity), and homologous recombination has been previously shown to occur both naturally and, in the laboratory (14–18). To systematically verify this observation, all 84 isolated biofilm-defective transposon mutants were screened via a targeted PCR assay of the *gtfBC* locus. For this PCR we would expect a ∼9,400 bp fragment in strains where *gtfB* and *gtfC* were intact. However, this PCR yielded two fragments of ∼10,000 bp and ∼4,600 bp. Notably, this *gtfBC* double-banding has been shown in other studies (14, 19). Strains presumed to have undergone homologous recombination between *gtfB* and *gtfC* displayed a single ∼4,600 bp PCR product. Figure 5A shows the distinct banding patterns of the intact and recombined *gtfBC* loci. Out of the 84 biofilm-defective mutants screened, 20 strains showed evidence of the *gtfBC* recombination event by PCR (Figure 5A and Figures S3-S21), indicating a high frequency of this modification in the transposon mutant library population (estimated at 2×10^−2^ rate). Our observed variant frequency is close to the rate, ∼1 x 10^-3^, reported by Narisawa *et al*. (15). The 20 mutants with the defective *gtfBC* coding region were poor biofilm producers, consistent with the critical role of these glucosyltransferases in sucrose-dependent biofilm formation. These strains exhibited a significantly reduced biofilm phenotype, with an average Z-score of −3.8 (S.E. +/- 0.2) (Table S3). To characterize the nature of the recombination, all of the PCR-confirmed *gtfBC* recombination mutants were genome sequenced (Table S4). Analysis of these sequenced isolates showed the recombination event resulted in the loss of a DNA fragment averaging ∼4,600 bp. The exact size of the deleted segment varied among isolates, ranging from 3,197 bp to 5,485 bp, suggesting multiple independent or heterogeneous recombination events (Table S3). The location of the transposon insertions within the *gtfBC* recombination mutants were diverse, as shown in Table S4. In three instances the *gtfBC* recombination was in conjunction with a transposon insertion that might also impact sucrose-mediated biofilm formation: *gtfD* (07_F12), between SMU_909 and *gtfD* (55_E6), and *irvR* (90_H10).

**Figure 5.**
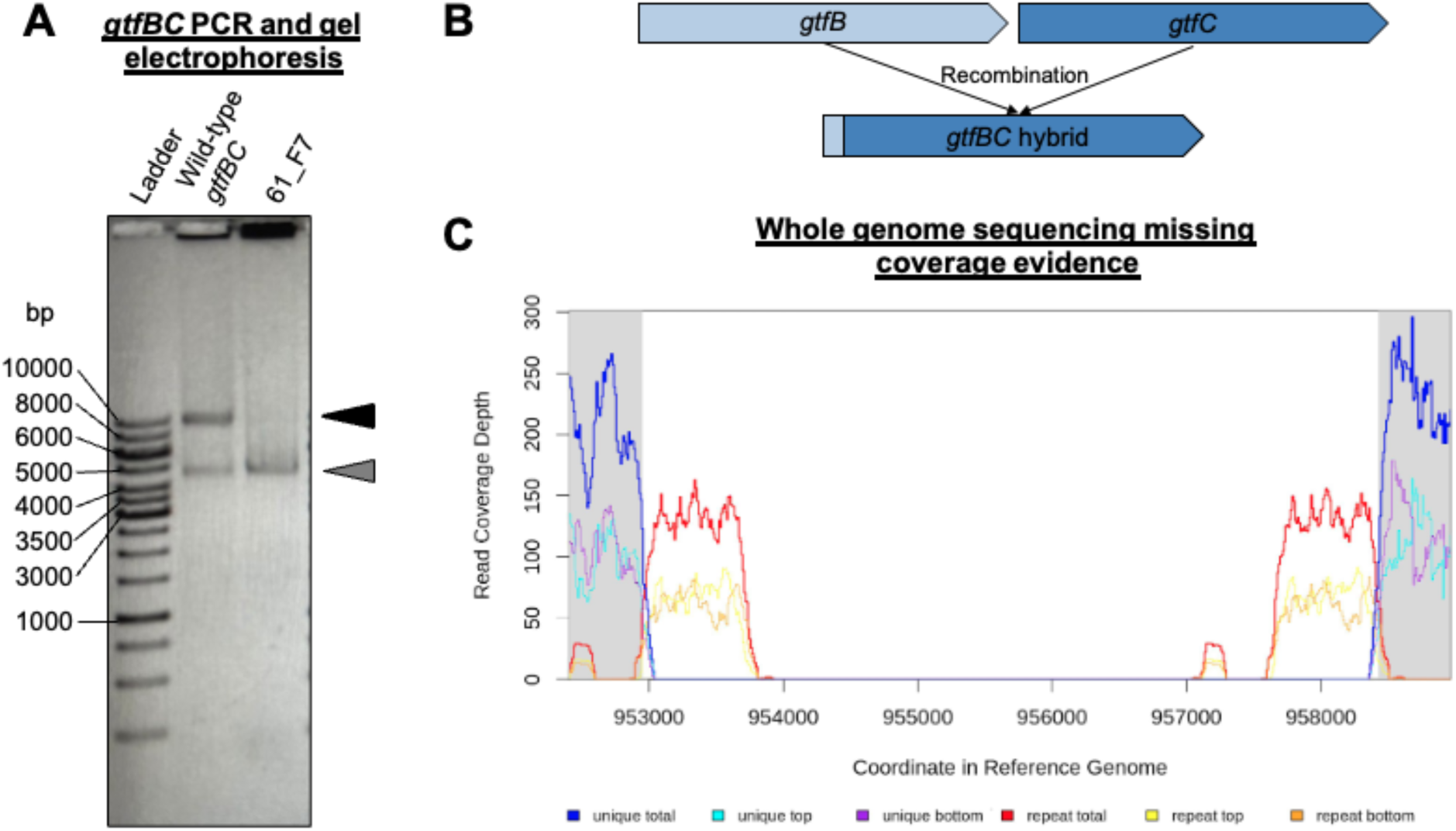
Molecular and genomic evidence of high-frequency recombination at the *gtfBC* locus. (A) To evaluate the genomic integrity of the highly homologous glucosyltransferase genes, PCR was performed using primers targeting the *gtfB* and *gtfC* tandem arrangement. In the wild-type UA159 parent, two characteristic bands are observed at approximately 10 kbp and 4.6 kbp. In the biofilm-defective transposon mutant 61_F7, the absence of the larger 10 kbp product and the presence of a single ∼4.6 kbp band confirm a spontaneous large-scale deletion resulting from recombination within the locus. (B) A proposed model illustrating how homologous recombination between the conserved sequences of *gtfB* and *gtfC* results in the excision of the intervening DNA and the formation of a single, hybrid glucosyltransferase gene. This rearrangement accounts for the ∼4.6 kbp deletion observed in biofilm-deficient variants. (C) Whole-genome sequencing coverage map of mutant 61_F7. Mapping of sequence reads reveals a definitive lack of coverage across the *gtfBC* genomic coordinates. The absence of read depth in this region provides physical confirmation of the deletion identified by PCR, illustrating that the severe biofilm-deficient phenotype in this strain is a consequence of genomic rearrangement rather than the primary transposon insertion.

Genome sequencing of the 84 transposon mutants revealed an unexpectedly high frequency of loss of the integrative and conjugative element (ICE) Tn*Smu1* (Figure 6). This element was absent in six of the 84 sequenced strains (57_D1, 67_D11, 67_E5, 76_D9, 92_C9, and 92_B7), representing a 7.1% loss frequency within the genome sequenced population. This observed rate is approximately 1,000-fold higher than the previously reported 0.006% spontaneous excision rate for *S. mutans* in planktonic cultures (20, 21). To determine if this high-frequency loss of Tn*Smu1* could account for biofilm-deficient phenotypes observed in the library, a targeted ΔTn*Smu1* deletion mutant was constructed. In sucrose-mediated biofilm assays, the ΔTn*Smu1* strain showed no statistically significant difference in biofilm formation compared to the wild-type UA159 parent strain (Figure 6B). These data confirm that the loss of TnSmu1 itself does not cause a biofilm defect and instead appears to occur spontaneously in our mutant library.

**Figure 6.**
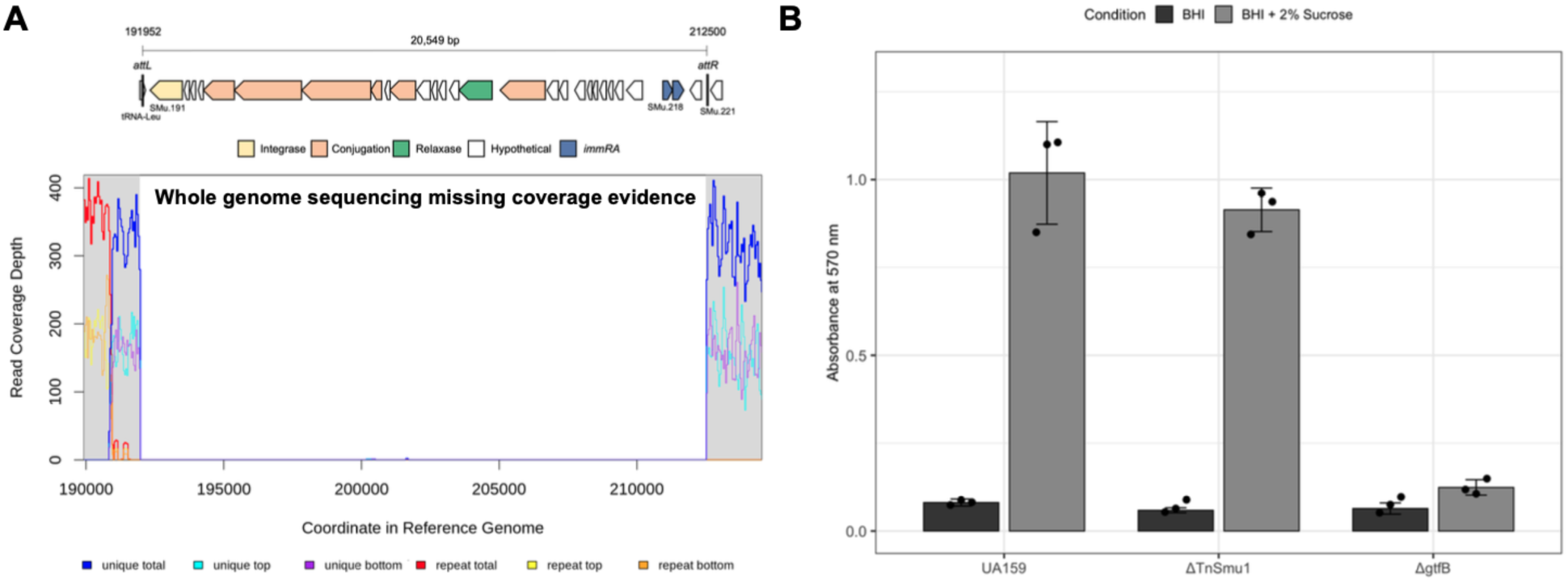
Validation of Tn*Smu1* genomic excision and its impact on biofilm formation. (A) Whole-genome sequencing read depth is displayed for a representative transposon mutant, illustrating the characteristic absence of coverage across the approximately 20 kbp integrative and conjugative element (ICE) known as Tn*Smu1*. This lack of reads confirms the spontaneous excision of the element from the chromosome during library construction or screening. (B) Quantitative assessment of biofilm biomass for *S. mutans* UA159, a targeted Tn*Smu1* deletion strain (ΔTn*Smu1*; LKM68), and a *gtfB* deletion strain (Δ*gtfB*). Biofilm formation was measured via crystal violet staining following growth in media supplemented with either glucose or sucrose. While the Δ*gtfB* strain exhibits a severe and significant defect specifically in sucrose-supplemented media, no statistical difference was observed between the wild-type UA159 parent and the ΔTn*Smu1* strain, confirming that the spontaneous loss of the ICE does not contribute to the biofilm-deficient phenotypes identified in the screen.

Lastly, the mutation annotation pipeline also identified multiple instances of missing coverage evidence for insertion sequence (IS) elements. Specifically, the breseq software identified that the ISSmu1 IS elements at SMU_436c and SMU_565c-SMU_566c were deleted. However, on further analysis we believe that the missing coverage evidence is ambiguous. This is because there are six copies of ISSmu1 in UA159 (22, 23) that share repetitive sequences, and because we were using short read sequencing to identify genomic variation, we are unable to identify these regions accurately.

### Characterization of newly identified biofilm determinants

To validate the functional relevance of novel hits identified in the arrayed screen, we generated targeted deletion mutants for three previously uncharacterized genes: SMU_635, SMU_1803c, and SMU_2160. To ensure the genomic integrity of the constructed mutants, we performed whole-genome sequencing on the ΔSMU_635, ΔSMU_1803c, and ΔSMU_2160 strains; comparative analysis confirmed that each mutant contained only the intended deletion.. We then compared their sucrose-dependent biofilm phenotypes to the wild-type UA159 parent (A570 = 0.402 ± 0.015) and a biofilm-deficient *ΔgtfB* control strain (A570 = 0.216 ± 0.044; p = 0.02) (Figure 7).

**Figure 7.**
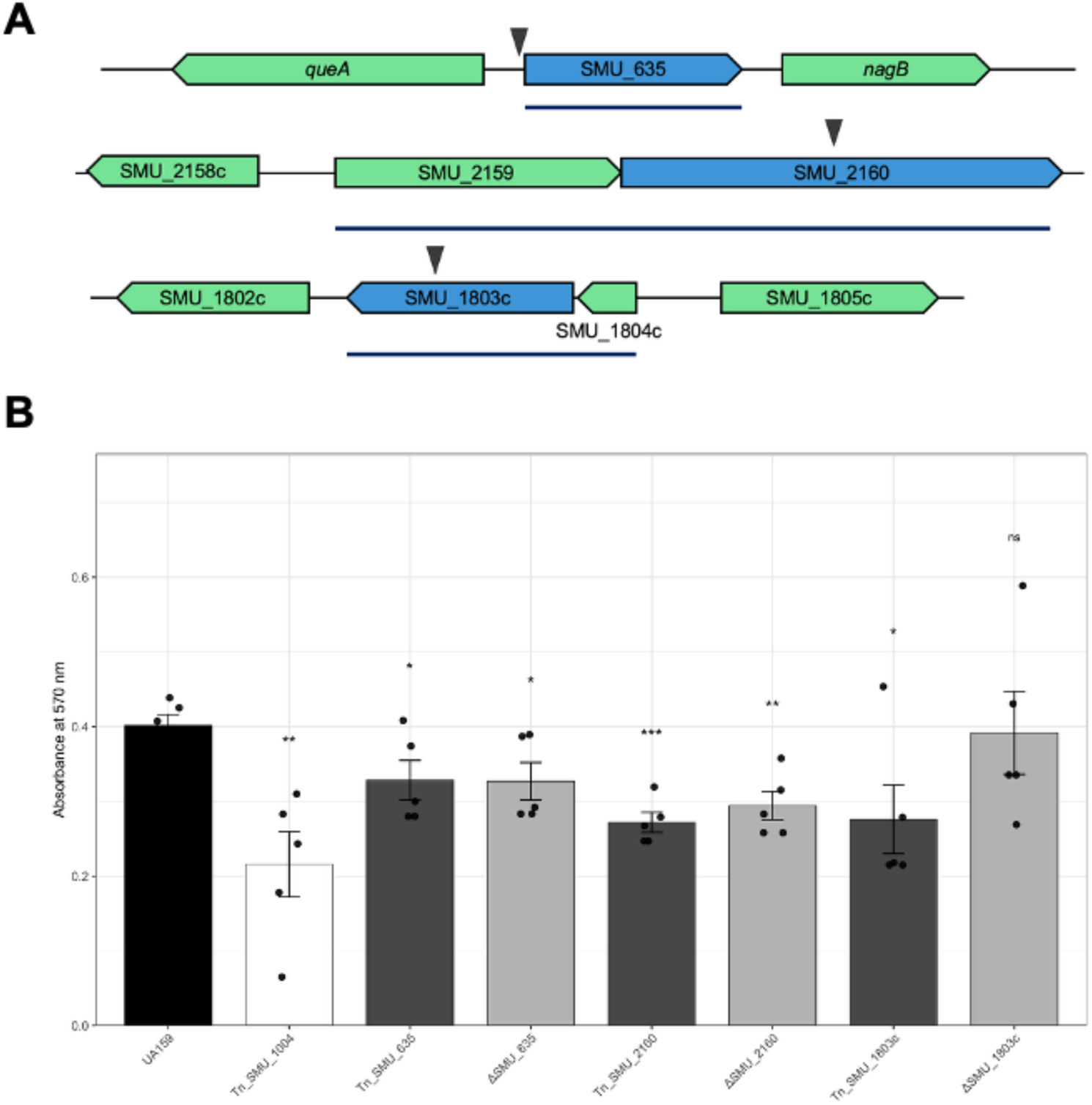
Genomic context and validation of novel sucrose-dependent biofilm determinants. (A) Genetic neighborhood of three novel hits: SMU_635 (putative metal transporter), SMU_2160 (YfhO-family protein), and SMU_1803c (DUF4230-domain protein). The genomic context shows their proximity to other genes, such as *queA* and *nagB* flanking SMU_635. Triangles indicate the location of transposon insertions identified in the initial screen, and each blue line indicates putative operons as determined via RNAseq data. (B) Comparison of sucrose-mediated biofilm formation between wild-type UA159, the original transposon (Tn) mutants, and targeted deletion strains. A Δ*gtfB* (Tn_SMU_1004) strain served as a biofilm-deficient control. Biofilm biomass was measured via crystal violet staining. Targeted deletions of SMU_635 and SMU_2160 significantly reduced biofilm formation, successfully replicating the phenotypes of their respective transposon mutants. In contrast, the ΔSMU_1803c deletion strain did not exhibit a statistically significant defect compared to the wild-type, suggesting the initial screen result for this locus was a library-specific artifact. Error bars represent the standard error of at least three biological replicates.

SMU_635 is predicted to encode a membrane-embedded metal ion transporter belonging to the CCC1/VIT1 family as analysis with both BlastP and Foldseek identified a VIT1 domain (residues 27–235), characteristic of vacuolar iron transporters involved in cation homeostasis and detoxification. SeqHub predictions further suggest its role as a cation efflux pump, potentially crucial for survival in the oral environment where metal availability fluctuates. The gene is flanked by housekeeping metabolic enzymes, including S-adenosylmethionine:tRNA ribosyltransferase (*queA*) upstream and glucosamine-6-phosphate deaminase (*nagB*) downstream. Quantitative biofilm assays confirmed that the ΔSMU_635 deletion mutant exhibited a significant defect in sucrose-mediated biofilm formation (A570 = 0.327 ± 0.025; p = 0.011), replicating the phenotype observed in the initial transposon mutant (A570 = 0.328 ± 0.026; p = 0.014) (Figure 7B).

SMU_1803c was identified as a moderate biofilm-defective hit in the initial screen. This protein contains a DUF4230 domain and is predicted to be a small, membrane-associated factor potentially involved in ribosome biogenesis or quality control. Its genomic context strongly supports this, as it is co-oriented with three essential ribosome biogenesis factors: *yqeI*, *yqeH* (a 50S-binding GTPase), and *yqeG* (a phosphatase). Despite the strong ribosome-related context, the targeted deletion of SMU_1803c failed to produce a statistically significant biofilm defect compared to the wild-type strain (A570 = 0.392 ± 0.056; p = 0.418) (Figure 7B). This indicates that the defect observed in the arrayed library (A570 = 0.276 ± 0.046; p = 0.015) may have been a library-specific artifact or the result of polar effects on adjacent genes during the initial transposon screen.

The arrayed library screen identified SMU_2160 as a novel determinant of sucrose-mediated biofilm formation. Bioinformatic analysis using BlastP revealed that SMU_2160 encodes an 857-amino acid protein (∼100 kDa) containing a YfhO domain (residues 18–837). SMU_2160 is predicted to be located in the cell membrane with thirteen transmembrane domains and a large extracellular domain of ∼375 amino acids (Figure S22). The YfhO protein family is classified as a GT-C type glycosyltransferase. To verify the screening result, a targeted deletion mutant ΔSMU_2160 was constructed. Quantitative biofilm assays confirmed that the loss of SMU_2160 resulted in a significant reduction in biofilm biomass compared to the wild-type UA159 parent (A570 = 0.294 ± 0.019; p = 0.001), successfully recapitulating the phenotype of the original transposon mutant (OD570 = 0.272 ± 0.013; p = 0.001) (Figure 7B).

## Discussion

The establishment and systematic characterization of a sequence-defined arrayed transposon library represents a significant advancement in the functional genomics of *S. mutans* UA159. By pairing high-throughput phenotypic screening with CP-CSeq and comprehensive whole-genome verification, this study provides a dual-purpose framework: identifying novel metabolic and regulatory determinants of sucrose-mediated biofilm formation and rigorously assessing the genomic stability inherent to large-scale mutant collections. The findings reveal that while many biofilm-deficient phenotypes are linked to central biosynthetic and metabolic pathways, a substantial proportion is the result of a high-frequency recombination event at the *gtfBC* locus. In addition, although not related to any biofilm phenotype, there was significant spontaneous excision of the Tn*Smu1* integrative and conjugative element. These results emphasize that the utility of an arrayed library is fundamentally contingent upon the integrity of the underlying genetic background, establishing systematic genomic verification as an essential prerequisite for accurate gene-phenotype mapping in streptococcal research. This methodological framework ensures that identified hits reflect true biological requirements rather than library-construction artifacts, thereby increasing the reproducibility and precision of future high-throughput investigations.

A central goal of this study was to implement Cartesian Pooling-Coordinate Sequencing (CP-CSeq) to establish a sequence-defined resource. While the original description of this method by Vandewalle *et al*. (11) characterized it as a robust and straightforward approach for deconvoluting arrayed libraries, our experience suggests that achieving high theoretical efficiency in practice is a challenge. Out of the 9,216 arrayed mutants, we successfully assigned high-confidence genomic coordinates to 1,400 strains (15.2%), with an additional 431 mutants assigned at lower-confidence thresholds. This deconvolution rate is notably lower than those reported in other ordered collections, such as *Mycobacterium bovis* BCG at 77% (11), Klebsiella pneumoniae KPPR1 at 81% (24), *Escherichia coli* CFT073 at 64% (25), *Enterococcus faecalis* OG1RF at 61% (10), *Shewanella oneidensis* at 55% (26), and *Proteus mirabilis* HI4320 at 43% (12). Several factors likely contributed to reduced library diversity and subsequent mapping limitations in our final sequencing. Potential issues include overamplification of transposon libraries during PCR steps or the excessive loss of template DNA during pooling, extraction, and enzymatic processing. Furthermore, biological factors specific to streptococci, such as cell chaining or the inadvertent picking of multiple colonies into single wells, may have introduced combinatorial noise that obscured unique signatures. While multiple transposon insertions in a single genome could theoretically complicate deconvolution, previous Southern blot analyses suggest this is an unlikely primary cause for the observed rates (27). Together, the discrepancy between the ideal deconvolution rate and our observed results highlights several critical technical considerations. Future implementations might benefit from a probabilistic approach to recover low-confidence targets or a sheared DNA methodology to better preserve library diversity. Given these challenges, the most pragmatic takeaway is that this collection is best utilized as a traditional arrayed library, where high-throughput phenotypic screening is followed by targeted sequencing of interesting hits to confirm insertion sites and genomic backgrounds.

Regardless, our screen successfully validated the library’s utility by identifying a diverse set of biofilm-defective mutants. Our strategy of sampling strains across the entire defective range, rather than focusing only on the most severe phenotypes, proved effective. While the strongest defects (Z-score < −2.5) were overwhelmingly attributed to *gtfBC* or *gcrR* mutations, the moderate-defect group (Z-score −1 to −2.5) was significantly more genetically diverse. This moderate-defect group included numerous genes with established roles in *S. mutans* biology and biofilm formation, confirming the sensitivity of our high-throughput assay. These known genes can be broadly categorized into the following categories: (1) carbohydrate metabolism, (2) surface adhesion and matrix production, and (3) stress response regulation. Mutations in *scrB* (sucrose-6-phosphate hydrolase), *fruR*/*fruI* (fructose utilization), and *lacD2* (tagatose 6-phosphate aldolase) highlight the importance of diverse sugar processing pathways for biofilm maturation. We identified *brpA* (biofilm regulatory protein), *mubM* (mutanobactin biosynthesis), *pgfM1* (glycosyltransferase), and *rgpI* (glycosyltransferase), all of which are linked to cell wall integrity, adhesion, or eDNA release (28–32). Hits in the two-component system *ciaH* and ion transporters *mntH* (manganese) and *copA* (copper) align with their known roles in regulating stress responses and environmental adaptation crucial for biofilm development (33–35). The identification of these diverse sets of genes, all with previous links to biofilm adhesion or accumulation, provides critical confidence that our screening strategy and mutant library can robustly identify genes required for sucrose-mediated biofilm formation.

Beyond identified biofilm machinery, our unbiased sampling of the arrayed library enabled the discovery of genes not previously linked to sucrose-mediated biofilm formation in *S. mutans*. Specifically, targeted deletion mutants confirmed that SMU_635 and SMU_2160 are necessary for optimal biofilm growth, replicating the phenotypes observed in the original transposon screen. SMU_635 is predicted to encode a membrane-embedded metal ion transporter of the CCC1/VIT1 family, and putative regulatory analysis places it within the PerR regulon, which is a metal-dependent system that governs oxidative stress responses in *S. mutans* (36). Recent research showed significant down-regulation of SMU_635 in response to a halogenated phenazine antimicrobial (HP-29) that disrupts metal homeostasis (37). We anticipate that the disruption of SMU_635 may impair intracellular metal balance which is a critical factor for enzymatic function and oxidative stress tolerance during biofilm growth. SMU_2160 encodes a YfhO-family protein, a class of enzymes increasingly linked to the glycosylation of proteins and cell wall polymers in Gram-positive bacteria (32, 38). The *S. mutans* UA159 genome contains two other YfhO-domain homologs, SMU_2064c and SMU_2066c, which function as glycosyltransferases within the Pgf (protein glycosylation) system (39). Depending on the strain background, the Pgf system has been shown to glycosylate several *S. mutans* proteins, including those involved in biofilm and virulence mechanisms (31, 40). While further experiments will be required to confirm the specific substrate of SMU_2160, its genomic context, conserved across closely related streptococci, places it immediately downstream of a putative ATPase component of an ABC transporter. With this genomic arrangement we hypothesize that the SMU_2159/2160 complex facilitates the export and subsequent transfer of carbohydrate moieties onto surface-associated proteins or cell-wall polymers. Together, these results suggest that optimal biofilm development is not only dependent on dedicated adhesion machinery but is deeply integrated into fundamental aspects of cellular physiology, including metal-dependent stress responses and post-translational surface modifications.

A critical and unexpected outcome of our systematic screen was the high degree of genomic instability observed in the *S. mutans* UA159 mutant library. We identified two major classes of instability: homologous recombination at the *gtfBC* locus and excision of the ICE Tn*Smu1*. The most impactful event was spontaneous recombination between the highly homologous *gtfB* and *gtfC* genes, which occurred in 25% of the sequenced strains. This large-scale deletion, resulting in the loss of a ∼4.6 kb fragment, is consistent with previous reports (14–18), confirming this locus as a frequent hotspot for generating biofilm-deficient variants. A second, distinct instability event was the excision of the ICE Tn*Smu1*, which we detected in 7.1% of sequenced strains, a rate approximately 1,000-fold higher than the 0.006% spontaneous excision reported in planktonic cultures (20). This finding initially presented another major potential artifact. However, we conclusively demonstrated that this event was not a confounding factor for biofilm phenotypes, as a targeted ΔTn*Smu1* deletion mutant formed biofilms identically to the wild-type parent. Taken together, these two findings underscore that *S. mutans* can undergo significant genomic alterations during routine laboratory manipulation. Both Tn*Smu1* excision and *gtfBC* recombination are regulated processes known to be induced by cellular stress, such as DNA damage or growth on solid media (20, 21). It is probable that the stresses inherent to library construction and screening (e.g., *in vitro* transposition, growth on selective agar, or a part of the biofilm screen) provided an unknown signal(s) that induced these events. This serves as a critical methodological consideration for the field, demonstrating that genomic stability at mobile elements and repetitive regions cannot be assumed. Furthermore, it validates the necessity of whole-genome sequencing to accurately interpret screen results and differentiate true gene-phenotype links from high-frequency genomic artifacts.

This study provides a high-confidence genetic resource for *S. mutans* while establishing a new methodological standard for functional genomics. By identifying novel biofilm determinants such as SMU_635 and SMU_2160, we expand the known requirements for biofilm development. Crucially, the discovery of pervasive instability at the *gtfBC* locus and high-frequency Tn*Smu1* excision highlights a systemic risk in large-scale mutant screens: the misattribution of phenotypes due to undetected genomic rearrangements. Ultimately, our findings demonstrate that an arrayed library is only as defined as its underlying background, establishing systematic whole-genome verification as an essential prerequisite for accurate gene-phenotype mapping in streptococcal research.

## Materials and Methods

### Bacterial strains and growth conditions

*S. mutans* strains were routinely cultured in Brain Heart Infusion (BHI) broth (BD Difco) at 37°C in a 5% CO_2_ environment. Antibiotics were used at the following concentrations: 1 mg/mL spectinomycin and 1 mg/mL kanamycin. A list of strains (Table S5) can be found in the supplementary material.

### Construction of the arrayed mutant library

*S. mutans* UA159 Tn mutants library generated by Shields et al. (2018) stored at –80 °C were diluted at 10^-6^ and incubated in a 5% CO2 environment on BHI agar at 37 °C for 48 h. After incubation, 9,216 single Tn-mutants were picked from BHI-spectinomycin agar using sterile toothpicks and inoculated into 96-deep-well plates with BHI-spectinomycin media. Plates were incubated at 37°C in a 5% CO_2_ environment. Subsequently, 500 µL were taken from the deep-well plates and 50% glycerol was added 1:1 to each well before storage at –80 °C. Each arrayed plate was replicated three times and assigned a unique identifier (ex. Tn-SmUA159-P1).

### Biofilm-defective mutant screening

Transposon mutant library plates were removed from the cryofreezer to thaw at room temperature. During this time, a ninety-six well microtiter plate was prepared containing 100 µL of BHI containing 2% sucrose (w/v). Adding sucrose to the BHI solution allowed for the Tn mutants to express the sucrose-dependent biofilm phenotype for analysis. Transposon mutants from each arrayed mutant plate were then placed into the BHI-sucrose plate via a 48-pin replicator (Boekel Scientific). Microtiter plates were then incubated for 24 h at 37°C in a 5% CO_2_ incubator. The following day the microtiter plates were removed from the incubator and images of the biofilm growth were captured using a microplate reader. Any individual wells that had no visual planktonic growth were noted to be exempt from further screenings, since a lack of growth would explain any reduced biofilm formation. Following this, all media within the biofilm plate wells was discarded into media waste. Distilled water (dH_2_O) was then used to wash loosely adherent cells from each well. One hundred microliter of dH_2_O was pipetted into each well and discarded for three total repetitions. Next, one hundred µL of 0.05% (w/v) crystal violet (CV) was then added to each well to stain for approximately 15 min at room temperature. The staining solution was then discarded, and the wells were washed with dH_2_O three times. Next, 100 µL of acetic acid (7 % v/v) was then added into the wells to resuspend the CV into solution. The absorbance of the biofilm stain was measured at 570 nm using a microplate reader (Synergy HT), and results were recorded in Microsoft Excel. Photographic images of the plate were also taken to compare to the images of the biofilm before the staining.

To determine if a mutant strain had significantly different biofilm formation compared to wild-type we used a standardized Z-score method. Using this method, a negative z-score can indicate a reduced ability to form a biofilm, while a positive z-score can indicate an increased ability. First, to account for variations in biofilm growth between biological replicates, we calculated the average CV absorbance reading for each transposon mutant plate; with the assumption that virtually all mutants would display a wild-type biofilm forming ability. Then, to calculate the Z-score for each transposon mutant, we used the following formula: (individual mutant well absorbance - plate average well absorbance)/standard deviation of the plate average well absorbance. Average Z-scores were calculated for each mutant well position from three biological replicates. We selected biofilm-defective mutants on the basis of a < −1 Z-score (< 25% biofilm formation compared to wild-type) and no visual defects in planktonic growth. After selecting 84 transposon mutants, they were individually cultured and stored at –80 °C and verification growth and biofilm assays were completed.

### Transposon mutant pooling for CP-CSeq

The location of the Tn insertion in each of the mutants within the library was determined using CP-CSeq strategy, which combines combinatorial pooling and next-generation Illumina sequencing to reduce the number of sequencing samples from 9,216 to 40, with each mutant represented in three distinct groups (11). The pool consisted of combining all mutants (1) in the same rows (25 µL of each well; 8 samples; X column pools), (2) in the same columns (50 µL of each well; 12 samples; Y column pools), and (3) from a single plate into one well of a 96-well plate, followed by the pooling of the rows and columns, like steps 1 and 2 (100µL of each well for Y column pools; 150µL of each well for X column pools; 20 samples; Figure S1). After the mutants were pooled, the samples were placed at −80 °C for storage.

### Transposon sequencing library preparation

Genomic DNA (gDNA) was extracted from each of the 40 combinatorially pooled mutant samples using the MasterPure™ Gram Positive DNA Purification Kit (BioSearch Technologies), following an optimized protocol (Walker & Shields, 2022). Briefly, overnight cultures were pelleted and resuspended in 150 µL of spheroplasting buffer. Cell wall digestion was performed with 10 µL of lysozyme (25 mg/mL; MP Biomedicals, USA) at 37°C, followed by addition of 150 µL of lysis solution and disruption with 0.1 mm diameter glass beads using a Mini-Beadbeater-24 (BioSpec) with 3 cycles of 30 s with ice incubation between cycles. Proteinase K (1 µL) was added to each sample and incubated at 65 °C with intermittent vortexing. After cooling, RNase A (1 µL) was added and samples were incubated at 37°C. Lysates were cleaned using 175 µL of MPC Protein Precipitation Reagent, and gDNA was precipitated with isopropanol, washed with 75% ethanol, air-dried, and resuspended in 25 µL of TE buffer. DNA concentrations were quantified using the Qubit dsDNA BR Assay Kit and Qubit Fluorometer (Invitrogen), following the manufacturer’s instructions. Next, gDNA was digested with MmeI (New England Biolabs) for 4 h at 37°C. Digested samples were purified by phenol:chloroform:isoamyl alcohol extraction (25:24:1; Thermo Scientific), followed by ethanol precipitation and washes with 70% ethanol. Pellets were air-dried, and DNA was resuspended in 25 µl of dH_2_O. Barcode-containing adapter ligation was performed overnight at 16 °C using T4 DNA ligase (400 U/µl; New England Biolabs). After ligation, transposon regions were PCR-amplified and products were separated on 1.8% agarose gels stained with SYBR Safe Gel Stain (10,000X; Invitrogen). Bands of ∼120 bp were excised, purified using QIAquick Gel Extraction Kit (QIAGEN), and eluted in 50 µL dH_2_O. DNA yields were determined using Qubit Fluorometer (Invitrogen), and samples were pooled into two 1.5 mL microcentrifuge tubes for sequencing. Oligonucleotides were ordered from Integrated DNA Technologies (IDT; Table S6).

### Illumina sequencing

Pooled libraries were normalized to 5 nM and sequenced at SeqCenter (Pittsburgh, USA) using Illumina NovaSeq with 150 cycle SP kit (2 x 75 bp paired end reads). To compensate for low sequence diversity characteristic of Tn-seq libraries, a 50% PhiX spike-in (Illumina Control Library) was included to enhance base-calling accuracy and calibration. This sequencing setup ensures sufficient read length to capture transposon-flanking regions while maintaining high-quality data for precise insertion mapping.

### Validation of CP-CSeq deconvolution

Sequence data processing and CP-CSeq were conducted on the HiPerGator high-performance computing cluster (University of Florida), running a Linux environment with Bash scripting (University of Florida Research Computing, n.d.). Scripts used for processing are provided in the supplemental materials. Illumina reads were demultiplexed by barcode, and the 16 bp downstream of each transposon insertion site were extracted. Reads were aligned to the *S. mutans* UA159 reference genome using Burrows-Wheeler Aligner (BWA) aln algorithm optimized for short reads (Li & Durbin, 2009). Sorted BAM files were processed using HTSeq to quantify reads mapped to genomic features (Anders et al., 2015).

### Validation of CP-CSeq deconvolution

For validation, 20 transposon mutants were randomly selected from the library and subjected to PCR to confirm the accuracy of the transposon insertion CP-CSeq deconvolution. Gene sequences were retrieved from BV-BRC, and primers targeting transposon and gene-specific regions were designed using Primer3Plus. Mutants were cultured on BHI plates containing spectinomycin at 37°C in a 5% CO_2_ environment for 48 hours. After incubation, single colonies were picked with sterile toothpicks and used as templates for PCR amplification with their corresponding set of primers (Table S6). PCR products were separated on 1.8% agarose gels stained with SYBR Safe Gel Stain.

### Identification of *gtfBC* variants

A PCR-based assay was used to determine if a homologous recombination event had occurred between the *gtfB* and *gtfC* genes for each transposon mutant. Forward (*gtfBC*_F; tcagttttaagaggacggcac) and reverse (*gtfBC*_R: agcctgagaaatttacagctca) primers were designed so that they would amplify the entire *gtfBC* region. LongAmp Taq 2X Master Mix (New England Biolabs) was combined with the *gtfBC* primers, 8.5 µL of nuclease-free water, and 1.5 µL of the extracted *S. mutans* DNA. PCR was performed under the following conditions: an initial denaturation at 94 °C for 5 minutes, followed by 35 cycles consisting of denaturation at 94 °C for 10 seconds, annealing at 56 °C for 30 seconds, and extension at 65 °C for 10 minutes. This was followed by a final extension at 65 °C for 10 minutes, with a final hold at 4 °C. *gtfBC* fragments were visualized using agarose gel electrophoresis.

### Biofilm loss-of-function mutation identification

Transposon mutant gDNA was extracted using the MasterPure™ Gram Positive DNA Purification Kit as described above, for 84 biofilm-defective mutants. Transposon mutant DNA samples were sent to SeqCenter for Illumina whole genome sequencing. We selected the 200 Mbp sequencing package for bacterial genomes <5 Mbp with haploid variant calling. For the variant calling, transposon mutant genomic information was compared to the reference *S. mutans* UA159 genome (AE014133.2) using the Breseq software package (41). This software identifies single base mutations, insertions and deletions, and chromosome rearrangements. For each mutant, we used the fastq sequencing files to identify the location of the transposon insertion by searching for the inverted repeat sequence (TAACAGGTTGGATGATAAGT) and using blastn to identify the region upstream of the inverted repeat in the *S. mutans* UA159 genome. To discover genome variants we used the breseq data file, called index.html, to identify differences between our laboratory-adapted version of UA159 versus each transposon mutant strain. Variant information, and the transposon insertion location, were recorded in a database for each of the sequenced transposon mutants.

### Gene mutagenesis

Targeted gene deletions were generated to validate phenotypes from the transposon screen by replacing selected genes (Table S4) with the *aphA3* cassette using PCR ligation mutagenesis (42). Gene sequences were retrieved from BV-BRC, and 400-600 bp upstream and downstream homologous arms were designed using Primer3Plus. Flanking regions were amplified from *S. mutans* UA159 genomic DNA, and the *aphA3* cassette was obtained from plasmid pALH123 isolated from *E. coli* using the QIAprep Miniprep Kit (QIAGEN). PCR products and plasmid were digested for 1 h at 37 °C, purified using the QIAquick Gel Extraction Kit, and ligated together using T4 DNA ligase for 1 h at room temperature. Ligation mixtures were transformed into competent *S. mutans*, followed by selection on BHI-kanamycin plates at 37°C in a 5% CO_2_ environment for 48 hours. Colonies were screened by PCR, and products were separated on 1.8% agarose gels stained with SYBR Safe Gel Stain. Verified mutants underwent genomic DNA extraction and whole-genome sequencing at SeqCenter.

## Acknowledgements

This work was supported by the National Institute of Dental and Craniofacial Research (NIDCR) grants DE034550 (MAA) and DE033403 (RCS and LKM). This work was also supported by the Arkansas INBRE program [NIGMS P20 GM103429] (RCS and MAA).

## References

1. Lemos J, Palmer S, Zeng L, Wen Z, Kajfasz J, Freires I, Abranches J, Brady L. 2019. The Biology of *Streptococcus mutans*. Microbiol Spectr 7:10.1128/microbiolspec.GPP3-0051–2018.

2. Bowen WH, Koo H. 2011. Biology of *Streptococcus mutans*-derived glucosyltransferases: role in extracellular matrix formation of cariogenic biofilms. Caries Res 45:69–86.

3. Matsumoto-Nakano M, Fujita K, Ooshima T. 2007. Comparison of glucan-binding proteins in cariogenicity of Streptococcus mutans. Oral Microbiol Immunol 22:30–35.

4. Lynch DJ, Fountain TL, Mazurkiewicz JE, Banas JA. 2007. Glucan-binding proteins are essential for shaping Streptococcus mutans biofilm architecture. FEMS Microbiol Lett 268:158–165.

5. Crowley PJ, Brady LJ, Michalek SM, Bleiweis AS. 1999. Virulence of a spaP Mutant ofStreptococcus mutans in a Gnotobiotic Rat Model. Infect Immun 67:1201–1206.

6. Love RM, McMillan MD, Jenkinson HF. 1997. Invasion of dentinal tubules by oral streptococci is associated with collagen recognition mediated by the antigen I/II family of polypeptides. Infect Immun 65:5157–5164.

7. Brady LJ, Maddocks SE, Larson MR, Forsgren N, Persson K, Deivanayagam CC, Jenkinson HF. 2010. The changing faces of Streptococcus antigen I/II polypeptide family adhesins. Mol Microbiol 77:276–286.

8. Shields RC, Zeng L, Culp DJ, Burne RA. 2018. Genomewide Identification of Essential Genes and Fitness Determinants of *Streptococcus mutans* UA159. mSphere 3:e00031–18.

9. Baba T, Ara T, Hasegawa M, Takai Y, Okumura Y, Baba M, Datsenko KA, Tomita M, Wanner BL, Mori H. 2006. Construction of *Escherichia coli* K-12 in-frame, single-gene knockout mutants: the Keio collection. Mol Syst Biol 2:2006.0008-2006.0008.

10. Dale JL, Beckman KB, Willett JLE, Nilson JL, Palani NP, Baller JA, Hauge A, Gohl DM, Erickson R, Manias DA, Sadowsky MJ, Dunny GM. 2018. Comprehensive Functional Analysis of the *Enterococcus faecalis* Core Genome Using an Ordered, Sequence-Defined Collection of Insertional Mutations in Strain OG1RF. mSystems 3:e00062–18.

11. Vandewalle K, Festjens N, Plets E, Vuylsteke M, Saeys Y, Callewaert N. 2015. Characterization of genome-wide ordered sequence-tagged Mycobacterium mutant libraries by Cartesian Pooling-Coordinate Sequencing. 1. Nat Commun 6:7106.

12. Pearson MM, Pahil S, Forsyth VS, Shea AE, Mobley HLT. 2022. Construction of an Ordered Transposon Library for Uropathogenic Proteus mirabilis HI4320. Microbiol Spectr 10:e03142–22.

13. Arjes HA, Sun J, Liu H, Nguyen TH, Culver RN, Celis AI, Walton SJ, Vasquez KS, Yu FB, Xue KS, Newton D, Zermeno R, Weglarz M, Deutschbauer A, Huang KC, Shiver AL. 2022. Construction and characterization of a genome-scale ordered mutant collection of Bacteroides thetaiotaomicron. BMC Biol 20:285.

14. Kimijima M, Narisawa N, Nakagawa-Nakamura T, Senpuku H. 2024. Allelic Variation in gtfB–gtfC Region of Natural Variant of Streptococcus mutans Without Biofilm Formation. Bacteria 3:369–378.

15. Narisawa N, Kawarai T, Suzuki N, Sato Y, Ochiai K, Ohnishi M, Watanabe H, Senpuku H. 2011. Competence-dependent endogenous DNA rearrangement and uptake of extracellular DNA give a natural variant of Streptococcus mutans without biofilm formation. J Bacteriol 193:5147–54.

16. Ueda S, Kuramitsu HK. 1988. Molecular basis for the spontaneous generation of colonization-defective mutants of Streptococcus mutans. Mol Microbiol 2:135–40.

17. Inagaki S, Fujita K, Takashima Y, Nagayama K, Ardin AC, Matsumi Y, Matsumoto-Nakano M. 2013. Regulation of Recombination between gtfB/gtfC Genes in Streptococcus mutans by Recombinase A. Sci World J 2013:405075.

18. Yamashita Y, Tomihisa K, Nakano Y, Shimazaki Y, Oho T, Koga T. 1999. Recombination between gtfB andgtfC Is Required for Survival of a dTDP-Rhamnose Synthesis-Deficient Mutant of Streptococcus mutans in the Presence of Sucrose. Infect Immun 67:3693–3697.

19. Hazlett KRO, Michalek SM, Banas JA. 1998. Inactivation of the gbpA Gene of Streptococcus mutans Increases Virulence and Promotes In Vivo Accumulation of Recombinations between the Glucosyltransferase B and C Genes. Infect Immun 66:2180–2185.

20. McLellan LK, Anderson ME, Grossman AD. 2022. TnSmu1 is a functional integrative and conjugative element in *Streptococcus mutans* that when expressed causes growth arrest of host bacteria. Mol Microbiol 118:652–669.

21. King S, Quick A, King K, Walker AR, Shields RCY 2022. Activation of TnSmu1, an integrative and conjugative element, by an ImmR-like transcriptional regulator in *Streptococcus mutans*. Microbiology 168:001254.

22. Old LA, Russell RRB. 2008. Distribution and activity of IS elements in Streptococcus mutans. FEMS Microbiol Lett 287:199–204.

23. Ajdić D, McShan WM, McLaughlin RE, Savić G, Chang J, Carson MB, Primeaux C, Tian R, Kenton S, Jia H, Lin S, Qian Y, Li S, Zhu H, Najar F, Lai H, White J, Roe BA, Ferretti JJ. 2002. Genome sequence of *Streptococcus mutans* UA159, a cariogenic dental pathogen. Proc Natl Acad Sci U S A 99:14434–9.

24. Mike LA, Stark AJ, Forsyth VS, Vornhagen J, Smith SN, Bachman MA, Mobley HLT. 2021. A systematic analysis of hypermucoviscosity and capsule reveals distinct and overlapping genes that impact Klebsiella pneumoniae fitness. PLOS Pathog 17:e1009376.

25. Shea AE, Marzoa J, Himpsl SD, Smith SN, Zhao L, Tran L, Mobley HLT. 2020. Escherichia coli CFT073 Fitness Factors during Urinary Tract Infection: Identification Using an Ordered Transposon Library. Appl Environ Microbiol 86:e00691–20.

26. Baym M, Shaket L, Anzai IA, Adesina O, Barstow B. 2016. Rapid construction of a whole-genome transposon insertion collection for Shewanella oneidensis by Knockout Sudoku. Nat Commun 7:13270.

27. Shields RC, O’Brien G, Maricic N, Kesterson A, Grace M, Hagen SJ, Burne RA. 2018. Genome-wide screens reveal new gene products that influence genetic competence in *Streptococcus mutans*. J Bacteriol 200:16 e00508–17.

28. Rainey K, Michalek SM, Wen ZT, Wu H. 2019. Glycosyltransferase-Mediated Biofilm Matrix Dynamics and Virulence of Streptococcus mutans. Appl Environ Microbiol 85:e02247–18.

29. Wen ZT, Baker HV, Burne RA. 2006. Influence of BrpA on Critical Virulence Attributes of *Streptococcus mutans*. J Bacteriol 188:2983.

30. Wu C, Cichewicz R, Li Y, Liu J, Roe B, Ferretti J, Merritt J, Qi F. 2010. Genomic Island TnSmu2 of Streptococcus mutans Harbors a Nonribosomal Peptide Synthetase-Polyketide Synthase Gene Cluster Responsible for the Biosynthesis of Pigments Involved in Oxygen and H2O2 Tolerance. Appl Environ Microbiol 76:5815–5826.

31. de Mojana di Cologna N, Andresen S, Samaddar S, Archer-Hartmann S, Rogers AM, Kajfasz JK, Ganguly T, Garcia BA, Saengpet I, Peterson AM, Azadi P, Szymanski CM, Lemos JA, Abranches J. 2024. Post-translational modification by the Pgf glycosylation machinery modulates Streptococcus mutans OMZ175 physiology and virulence. Mol Microbiol 122:133–151.

32. Rahman MM, Zamakhaeva S, Rush JS, Chaton CT, Kenner CW, Hla YM, Tsui H-CT, Uversky VN, Winkler ME, Korotkov KV, Korotkova N. 2025. Glycosylation of serine/threonine-rich intrinsically disordered regions of membrane-associated proteins in streptococci. Nat Commun 16:4011.

33. Qi F, Merritt J, Lux R, Shi W. 2004. Inactivation of the ciaH Gene in Streptococcus mutans Diminishes Mutacin Production and Competence Development, Alters Sucrose-Dependent Biofilm Formation, and Reduces Stress Tolerance. Infect Immun 72:4895–4899.

34. Kajfasz JK, Katrak C, Ganguly T, Vargas J, Wright L, Peters ZT, Spatafora GA, Abranches J, Lemos JA. 2020. Manganese Uptake, Mediated by SloABC and MntH, Is Essential for the Fitness of Streptococcus mutans. mSphere 5.

35. Garcia SS, Du Q, Wu H. 2016. Streptococcus mutans Copper Chaperone, CopZ, is critical for biofilm formation and competitiveness. Mol Oral Microbiol 31:515–525.

36. Kajfasz JK, Zuber P, Ganguly T, Abranches J, Lemos JA. 2021. Increased Oxidative Stress Tolerance of a Spontaneously Occurring *perR* Gene Mutation in *Streptococcus mutans* UA159. J Bacteriol 203:e00535–20.

37. Kajfasz JK, Hosay HB, Gao Q, Huigens RW, Lemos JA. 2026. Zinc-enhanced activity of an antimicrobial halogenated phenazine against Streptococcus mutans and other gram-positive bacteria. mSphere 11:e00585–25.

38. Rismondo J, Percy MG, Gründling A. 2018. Discovery of genes required for lipoteichoic acid glycosylation predicts two distinct mechanisms for wall teichoic acid glycosylation. J Biol Chem 293:3293–3306.

39. Avilés-Reyes A, Freires IA, Besingi R, Purushotham S, Deivanayagam C, Brady LJ, Abranches J, Lemos JA. 2018. Characterization of the pgf operon involved in the posttranslational modification of Streptococcus mutans surface proteins. Sci Rep 8:4705.

40. Aviles-Reyes A, Miller JH, Simpson-Haidaris PJ, Hagen FK, Abranches J, Lemos JA. 2014. Modification of Streptococcus mutans Cnm by PgfS Contributes to Adhesion, Endothelial Cell Invasion, and Virulence. J Bacteriol 196:2789–2797.

41. Deatherage DE, Barrick JE. 2014. Identification of mutations in laboratory evolved microbes from next-generation sequencing data using breseq. Methods Mol Biol Clifton NJ 1151:165–188.

42. Lau PCY, Sung CK, Lee JH, Morrison DA, Cvitkovitch DG. 2002. PCR ligation mutagenesis in transformable streptococci: application and efficiency. J Microbiol Methods 49:193–205.

